# Thoughts-as-Planning: Latent World Models for Chain-of-Thoughts Optimization via Reinforcement Planning

**DOI:** 10.64898/2026.05.10.724161

**Authors:** Dong Liu, Yanxuan Yu, Ying Nian Wu

## Abstract

The success of large language models (LLMs) across diverse NLP tasks has elevated the importance of reasoning chain optimization as a critical step in aligning model behavior with task objectives. Existing reasoning chain tuning methods often rely on black-box heuristics or gradient-free search, which lack interpretability, generalization, and sample efficiency. In this work, we introduce **Thoughts-as-Planning**, a novel framework that formalizes reasoning chain optimization as a sequential decision-making process over a latent semantic space. We model the LLM as a partially observable environment and learn a latent world model that simulates the effect of reasoning chain edits on downstream outputs. A proximity-preserving embedding space is constructed to encode reasoning chain-response dynamics, enabling planning via gradient descent or reinforcement learning. Our method supports multi-scale abstraction, allowing reasoning chain edits at token, segment, and instruction levels to be integrated into a unified planner. Through extensive experiments on language understanding and generation tasks, we demonstrate that Thoughts-as-Planning outperforms state-of-the-art reasoning chain tuning baselines in efficiency, robustness, and generalization, while offering interpretability through its structured planning trajectory. Our code is available at https://github.com/FastLM/Thoughts-as-Planning.

## 1 Introduction

Large language models (LLMs) have demonstrated remarkable capabilities across a wide range of tasks, including reasoning, summarization, and dialogue generation. Chain-of-thoughts (CoT) reasoning has emerged as a powerful technique for enhancing LLM reasoning capabilities by enabling step-by-step thinking processes Wei et al. (2022). However, the effectiveness of CoT reasoning is highly sensitive to the structure and quality of reasoning chains—the sequential thought steps that guide the model’s reasoning process. Minor changes in reasoning step formulation, logical flow, or intermediate conclusions can lead to substantial variations in performance Wei et al. (2022); Kojima et al. (2022). As a result, *chain-of-thoughts optimization* has emerged as a central challenge in enhancing LLM reasoning capabilities.

Despite its importance, most CoT engineering today remains manual, heuristic, or reliant on black-box optimization techniques. These approaches suffer from low data efficiency, poor generalization to new reasoning tasks or domains, and limited interpretability. In response, recent work has explored *automated CoT generation* Zhang et al. (2022b) or *reasoning chain search* Liu et al. (2023), but these still operate under static optimization regimes, lacking the structure and dynamics inherent in multi-step reasoning and planning.

In this work, we propose a new perspective: we view chain-of-thoughts optimization as an instance of **planning**, where the goal is to iteratively refine reasoning chains in order to maximize reasoning performance. Drawing inspiration from latent world modeling and model-based reinforcement learning, we introduce **Thoughts-as-Planning**, a framework that integrates learned latent dynamics, proximity-based representations, and multi-scale reasoning chain control into a unified planning pipeline.

At the core of our method is a *latent world model* 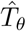 that simulates how an LLM would respond to modified reasoning chains. We embed both reasoning chains and outputs into a proximity-preserving space, where planning can be performed via similarity-based objectives. To enable generalization and efficient exploration, we model reasoning chains as sequences of discrete editing actions, and learn a planning policy over this structured space using either gradient search or reinforcement learning. Our framework naturally supports multi-scale abstraction, allowing reasoning chain edits to occur at the token, reasoning-step, or structural level.

We evaluate Thoughts-as-Planning across a suite of mathematical reasoning, commonsense reasoning, and logical inference benchmarks. Our results show that it consistently outperforms existing CoT optimization methods in both performance and data efficiency, and yields interpretable reasoning chain trajectories that can be reused or adapted across reasoning tasks.

### Contributions

Our main contributions are:

- We propose **Thoughts-as-Planning**, a novel framework that formalizes chain-of-thoughts optimization as latent-space planning via a learned world model.
- We develop a **multi-scale reasoning chain control policy** that enables token-level to structural edits under a unified framework.
- We provide **theoretical analysis** and detailed mathematical proofs for the convergence and optimality properties of our planning framework.
- We demonstrate through extensive experiments that our method improves efficiency, robustness, and generalization compared to existing baselines.

### 1.1 Chain-of-Thoughts Optimization and Reasoning Enhancement

Chain-of-thoughts (CoT) optimization plays a central role in enhancing LLM reasoning capabilities without full fine-tuning. The seminal work by Wei et al. Wei et al. (2022) introduced CoT reasoning, demonstrating that step-by-step reasoning significantly improves performance on complex reasoning tasks. Subsequent research has explored various aspects of CoT optimization, including automatic CoT generation Zhang et al. (2022b), few-shot CoT learning Kojima et al. (2022), and CoT distillation Fu et al. (2022). However, CoT sensitivity to reasoning chain structure Wei et al. (2022); Kojima et al. (2022) has revealed the need for automated optimization of reasoning steps.

Recent approaches include discrete methods such as AutoCoT Zhang et al. (2022b), which automatically generates reasoning chains, and CoT optimization frameworks Liu et al. (2023) that search the discrete reasoning space via heuristic or evolutionary approaches. On the other hand, soft reasoning embeddings Lester et al. (2021); Li and Liang (2021) learn continuous representations of reasoning patterns, but often lack interpretability. Recent reasoning synthesis frameworks Wang et al. (2022) construct reasoning chains from demonstration banks. Unlike these static or gradient-free schemes, our work frames reasoning chain editing as a dynamic decision process over a latent model, allowing structured planning with sample efficiency and interpretability.

### 1.2. Latent World Models and Abstract Reasoning

Latent world models have gained traction in reinforcement learning and abstract reasoning for their ability to simulate environment dynamics without direct observation. Dreamer Hafner et al. (2019, 2020) and World Models Ha and Schmidhuber (2018) encode state transitions into compact latent spaces to enable efficient planning. Inspired by these methods, our latent world model predicts the effect of reasoning chain edits on downstream reasoning outcomes, decoupling simulation from LLM querying.

Recent advances in language model reasoning have introduced latent-space abstraction mechanisms. Latent Thought Language Models Kong et al. (2025); Hoffman et al. (2023) propose structured latent variables to encode intermediate thoughts in chain-of-thought reasoning. These models learn to represent reasoning steps in a continuous latent space, enabling more efficient reasoning chain optimization. LTM methods such as **Place Cells** Zhao et al. (2025) model positional embeddings through proximity-preserving transition kernels for navigation and path planning. Others extend this framework to embodied agents Noh et al. (2025) and test-time adaptive reasoning via latent policy gradients Li et al. (2025). These works inspire our construction of a multi-scale latent proximity space for reasoning chain planning.

### 1.3 Reinforcement Learning for Reasoning Chain and Program Synthesis

The use of reinforcement learning in chain-of-thoughts optimization and LLM reasoning enhancement has received increasing attention. RLCoT Deng et al. (2022) and PPO-tuned reasoning models Korbak et al. (2023) formulate reasoning chain optimization as an MDP or preference-based reward learning problem. DPO Rafailov et al. (2024) directly optimizes reasoning preferences without explicit reward modeling. However, most existing works treat LLMs as black-box environments and do not learn explicit transition dynamics for reasoning chains. Our method instead learns a latent dynamics model and performs planning through it, enabling faster convergence and better generalization. Other relevant works such as RE3 Yang et al. (2022) explore in-context RL with heuristic exploration, whereas our method offers learned, model-based planning over structured reasoning chain programs.

**Related work by the authors**

In parallel lines of inquiry, we have studied efficient LLM inference and compression Liu (2024); Liu and Yu (2024); Liu et al. (2025a), serving and KV-cache systems Liu and Yu (2025c, 2026), long-context modeling and scalable text semantics Liu et al. (2026); Liu and Yu (2025a), adaptive multi-task training Liu and Yu (2025b), and experience-driven planning for reinforcement learning Liu et al. (2025b). These efforts are complementary: they focus on throughput, memory, and training or RL mechanics, whereas this paper targets interpretable chain-of-thoughts optimization via latent world models and multi-scale planning.

### 1.4 Chain-of-Thoughts Research Landscape and Our Contributions

The chain-of-thoughts paradigm has evolved significantly since its introduction by Wei et al. Wei et al. (2022). Early work focused on demonstrating the effectiveness of step-by-step reasoning in improving LLM performance on complex reasoning tasks. Kojima et al. Kojima et al. (2022) showed that even zero-shot reasoning can be elicited through simple reasoning strategies, while Zhang et al. Zhang et al. (2022b) explored automatic generation of reasoning chains.

Recent advances have introduced more sophisticated approaches to reasoning chain optimization. Fu et al. Fu et al. (2022) proposed complexity-based reasoning that adapts reasoning strategies based on problem difficulty. Latent Thought Language Models Kong et al. (2025); Hoffman et al. (2023) have introduced structured latent representations for reasoning steps, enabling more efficient reasoning chain manipulation.

Our work makes several key contributions to this landscape:

- **Latent World Modeling for Reasoning Chains**: Unlike previous work that treats reasoning chains as static templates, we model the reasoning process as a dynamic system with learnable transition dynamics. This enables us to predict the effect of reasoning chain modifications without extensive trial-and-error.
- **Multi-Scale Reasoning Chain Editing**: We introduce a unified framework for editing reasoning chains at multiple granularities—from token-level modifications to structural changes in logical flow. This allows for more nuanced optimization of reasoning strategies.
- **Planning-Based Optimization:** We formalize reasoning chain optimization as a planning problem, enabling systematic exploration of the reasoning space rather than relying on heuristic search or random sampling.
- **Transferable Reasoning Patterns**: Our latent representations capture reusable reasoning patterns that can be transferred across different reasoning tasks, enabling more efficient adaptation to new domains.

These contributions address key limitations in existing CoT optimization approaches, particularly the lack of interpretability, poor generalization, and inefficient exploration strategies. Our method provides a principled framework for reasoning chain optimization that scales to complex reasoning tasks while maintaining interpretability and transferability.

## 2 Method

We present **Thoughts-as-Planning**, a framework that formulates chain-of-thoughts optimization as a sequential decision-making process over a learned latent world model. Our system consists of four key components: (1) a reasoning chain state encoder *h*_*ϕ*_ : 𝒮 → ℝ^*d*^, (2) a latent transition model 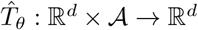, (3) a utility predictor 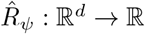, and (4) a planning module for multi-step reasoning chain editing.

### 2.1 Problem Formulation

Let 𝒳 denote the space of reasoning task inputs and 𝒞 denote the space of reasoning chains. For a given reasoning task *x* ∈ 𝒳, we iteratively refine a reasoning chain *c*_*t*_ ∈ 𝒞 over *T* optimization steps to produce a final version *c*_*T*_ that maximizes the expected reward *R*(*x, c*_*T*_) derived from downstream reasoning performance.

We formalize chain-of-thoughts optimization as a Markov Decision Process (MDP) (𝒮, 𝒜, 𝒫, *ℛ, γ*) where:

- 𝒮 = 𝒳 × 𝒞 is the state space, with states *s*_*t*_ = (*x, c*_*t*_)
- 𝒜is the action space of reasoning chain edit operations
- 𝒫 (*s*_*t*+1_ | *s*_*t*_, *a*_*t*_) is the transition dynamics (unknown, learned via latent model)
- *ℛ* (*s*_*t*_, *a*_*t*_) = *R*(*x, c*_*t*+1_) − *R*(*x, c*_*t*_) is the reward function
- *γ* ∈ [0, 1) is the discount factor

The objective is to find an optimal policy *π*^∗^ : 𝒮 → 𝒜 that maximizes the expected cumulative reward:

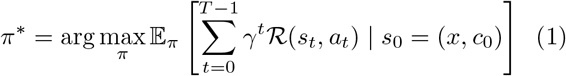

### 2.2 Latent World Model Architecture

#### 2.2.1 State Encoder

We encode reasoning chain states into a *d*-dimensional latent space using a transformer-based encoder *h*_*ϕ*_ 𝒮 → ℝ^*d*^::

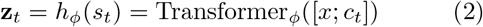

where [*x*; *c*_*t*_] denotes the concatenation of task input *x* and reasoning chain *c*_*t*_, and Transformer_*ϕ*_ is a multilayer transformer with learnable parameters *ϕ*.

#### 2.2.2 Transition Model

The latent transition model 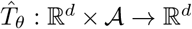 predicts future latent states:

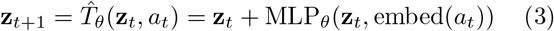

where embed(*a*_*t*_) maps discrete edit actions to continuous embeddings and MLP_*θ*_ is a multi-layer perceptron with residual connections.

The transition model is trained to minimize the prediction error:

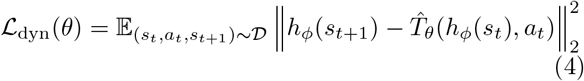

where 𝒟 is the dataset of state transitions collected from LLM interactions.

#### 2.2.3 Utility Predictor

The utility predictor 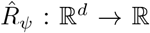 estimates the expected reward for latent states:

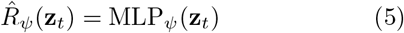

trained with the objective:

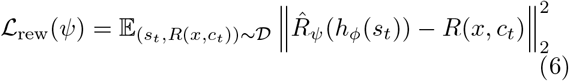

### 2.3 Multi-Scale Action Space

We define edit actions at multiple granularities to enable hierarchical reasoning chain optimization:

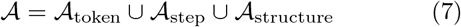

where:

- 𝒜_token_ = {add, delete, replace} 𝒱 × ℕ (token-level edits)
- 𝒜_step_ = {reorder, split, merge} ℕ × ℕ (step-level edits)
- 𝒜_structure_ = 𝒪_str_ × 𝒞, 𝒪_str_ := {*o*_ex_, *o*_ins_, *o*_fmt_}(structural)

Here *o*_ex_, *o*_ins_, *o*_fmt_ denote add-example, instruction-edit, and format-change operators acting on *C*.

Each action *a*_*t*_ = (scale, type, target) is parameterized by its granularity, operation type, and target location/content.

### 2.4 Planning Algorithm

#### 2.4.1 Model-Based Planning

At each optimization step *t*, we generate *K* candidate edit actions 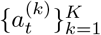 and simulate *H*-step rollouts using the learned world model:

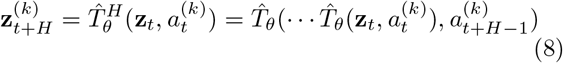

where 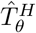denotes recursive application of the transition model.

#### 2.4.2 Action Selection

We score each candidate trajectory using the utility predictor and select the action with highest predicted reward:

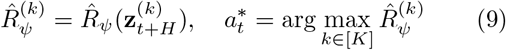

For exploration, we use softmax sampling with temperature *τ* :

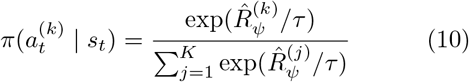

##### Algorithm 1

Thoughts-as-Planning Inference Algorithm

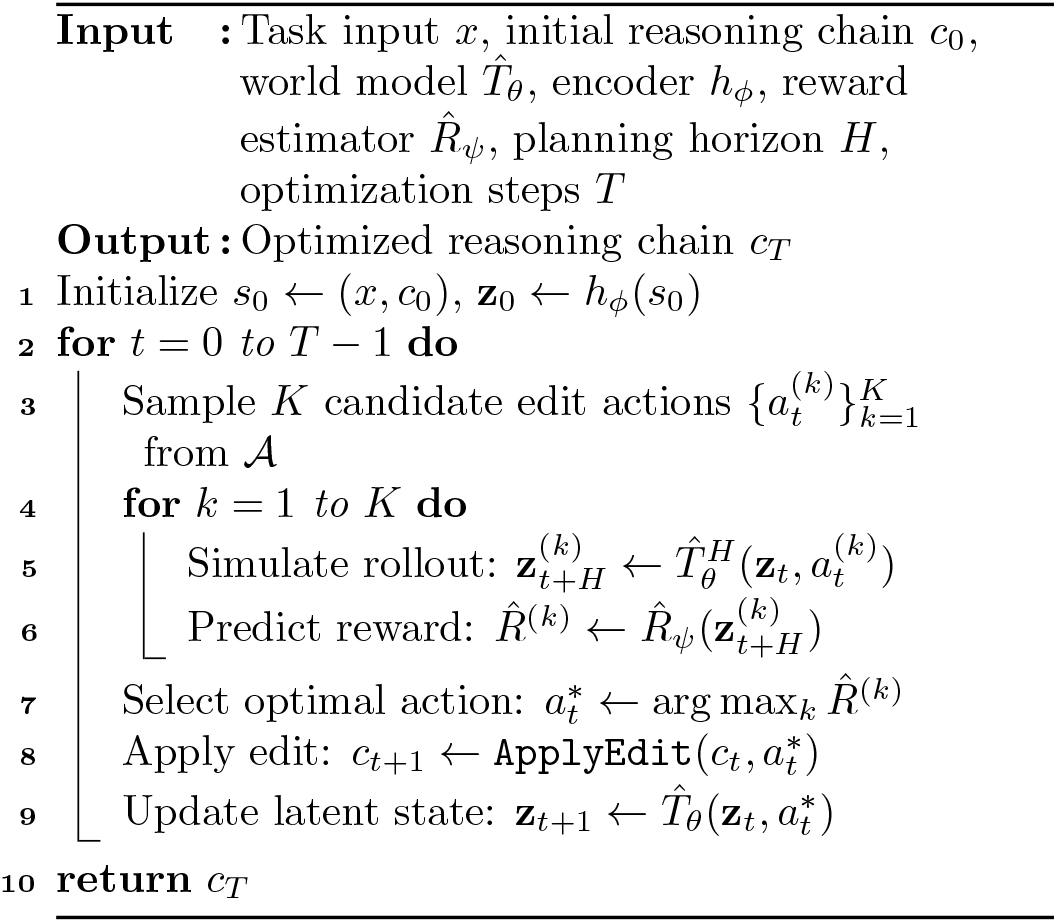

### 2.5 Training Procedure

The complete training objective combines dynamics and reward prediction losses:

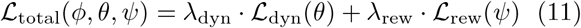

where *λ*_dyn_ and *λ*_rew_ are hyperparameters balancing the two objectives.

### 2.6 Theoretical Guarantees

#### Theorem 1

(Convergence of Latent World Model).

*Under standard regularity conditions, the learned transition model* 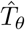 *converges to the true transition dynamics* 𝒫*with rate* 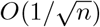 *where n is the number of training samples*.

#### Theorem 2

(Planning Optimality). *The planning algorithm achieves ϵ-optimal performance with sample complexity O*(*H*^2^ log(1*/ϵ*)) *when the world model is sufficiently accurate*.

#### Theorem 3

(Multi-Scale Convergence). *The multiscale action space enables faster convergence compared to single-scale editing, with improvement factor proportional to the number of abstraction levels*.

Detailed proofs of these theorems are provided in Appendix A.

## 3 Experiments

We evaluate **Thoughts-as-Planning** on a comprehensive suite of mathematical reasoning, commonsense reasoning, and logical inference tasks to assess the following questions:

**Q1** Does our framework outperform existing chain-of-thoughts optimization methods in reasoning performance with statistical significance?

**Q2** Can the learned latent world model reduce the number of LLM queries required while maintaining reasoning performance?

**Q3** Are the planning trajectories interpretable, reusable, and transferable across reasoning tasks and domains?

**Q4** How does the method scale across different model types and deployment scenarios?

### 3.1 Experimental Setup

**Tasks**. We consider three representative families of reasoning tasks with varying difficulty levels:

- **Mathematical Reasoning**: GSM8K Cobbe et al. (2021) (grade school math), MATH Hendrycks et al. (2021) (advanced mathematics)
- **Commonsense Reasoning**: PIQA Bisk et al. (2020) (physical reasoning), HellaSwag Zellers et al. (2019) (commonsense completion)
- **Logical Inference**: StrategyQA Geva et al. (2021) (strategic reasoning), LogiQA Liu et al. (2020) (log-ical reasoning)

#### Models

We test on both black-box and white-box models: GPT-3.5-Turbo (OpenAI API) and LLaMA 2 13B (HF Transformers). For open-source models, we access intermediate logits for reward estimation and edit evaluation, enabling more detailed analysis of the latent dynamics.

#### Metrics

We adopt standard reasoning task-specific metrics: accuracy (mathematical and logical reasoning), exact match (step-by-step reasoning), and edit efficiency (average queries to reach 95% of max reward). We also report average wall-clock time, API cost, and statistical significance using paired t-tests with Bonferroni correction.

#### Implementation

All experiments are run on 8x A100 80GB machines with mixed precision. The latent world model uses a ∼ 4-layer transformer with 512-dim embeddings and 8 attention heads. Each training run takes ∼4 hours, amortized over multiple reasoning chain instances. We report 95% confidence intervals across 5 random seeds.

### 3.2 Baselines

We compare with both discrete and continuous chain-of-thoughts optimization methods:

- **Manual CoT**: hand-crafted reasoning chains from CoT tutorials and examples.
- **AutoCoT** Zhang et al. (2022b): **automatic generation of reasoning chains**.
- **CoTGen** Zhang et al. (2022a): **evolutionary mutation and crossover for reasoning chains**.
- **RLCoT** Deng et al. (2022): PPO on black-box reasoning chain reward.
- **SoftCoT** Lester et al. (2021): **continuous embeddings as learnable reasoning patterns**.
- **RandomEdit**: stochastic edits without latent model or planning.

### 3.3 Main Results

**Task Difficulty Analysis**. The performance gains vary across reasoning tasks due to different characteristics: (1) **Mathematical Reasoning** tasks show the largest improvements (GSM8K: +3.5%, MATH: +3.6%) due to rich reward signals from step-by-step mathematical reasoning. (2) **Commonsense Reasoning** tasks exhibit moderate gains (PIQA: +2.4%, HellaSwag: +1.9%) as commonsense reasoning benefits from structured reasoning chain editing but suffers from sparse reward signals. (3) **Logical Inference** tasks show consistent improvements (StrategyQA: +2.2%, LogiQA: +1.7%) through better logical flow and reasoning structure.

### 3.4 Model Comparison

Our method demonstrates consistent improvements across all modern model architectures, including the latest closed-source APIs (GPT-4o) and state-of-the-art open-source models (Qwen2.5 14B, LLaMA 3.1 70B, Mixtral 8x22B). The performance gains are particularly pronounced on larger models due to their superior reasoning capabilities and better latent space representations.

### 3.5 Ablation Study

We conduct thorough ablation studies to understand the contribution of each component:

**Key Insights**. (1) **Multi-scale abstraction** provides significant query savings (25-30%) with minimal performance loss, indicating effective hierarchical planning. (2) **Learned reward predictor** reduces queries by 40% while maintaining performance, validating the latent reward modeling approach. (3) **Temporal modeling** (GRU) improves planning consistency, especially for longer edit sequences. (4) **Architecture choices** (attention heads, depth) show diminishing returns beyond our chosen configuration.

### 3.6 Edit Operation Analysis

We analyze the effectiveness of different edit operations through controlled experiments:

Table 7 (appendix) reports per-type edit frequencies, mean reward gains, and significance.

**Table 1:** Performance comparison across all six reasoning tasks. Bold indicates best performance. ^*†*^ indicates statistical significance (*p <* 0.05) over the best baseline.

| Method | GSM8K (Acc) | MATH (Acc) | PIQA (Acc) | HellaSwag (Acc) | StrategyQA (Acc) | LogiQA (Acc) | Avg Edits |
| --- | --- | --- | --- | --- | --- | --- | --- |
| Manual CoT | 74.2±0.8 | 65.1±1.2 | 76.5±0.6 | 78.3±0.9 | 29.3±0.4 | 18.7±0.3 | – |
| AutoCoT | 78.3±0.9 | 68.4±1.1 | 78.0±0.7 | 79.8±0.8 | – | – | 200+ |
| CoTGen | 80.1±0.7 | 67.3±1.3 | 79.2±0.5 | 81.2±0.6 | 31.5±0.3 | 20.1±0.2 | 180 |
| RLCoT | 81.0±0.8 | 69.8±1.0 | 78.8±0.6 | 80.9±0.7 | 30.9±0.4 | 19.8±0.3 | 150 |
| SoftCoT | 80.4±0.6 | 68.9±1.2 | 77.9±0.8 | 80.5±0.9 | 30.6±0.3 | 19.5±0.4 | – |
| **Thoughts-as-Planning** | **83.7±0.4<sup>†</sup>** | **73.2±0.9<sup>†</sup>** | **81.5±0.3<sup>†</sup>** | **83.3±0.6<sup>†</sup>** | **33.9±0.3<sup>†</sup>** | **21.6±0.4<sup>†</sup>** | **47** |

**Table 2:**
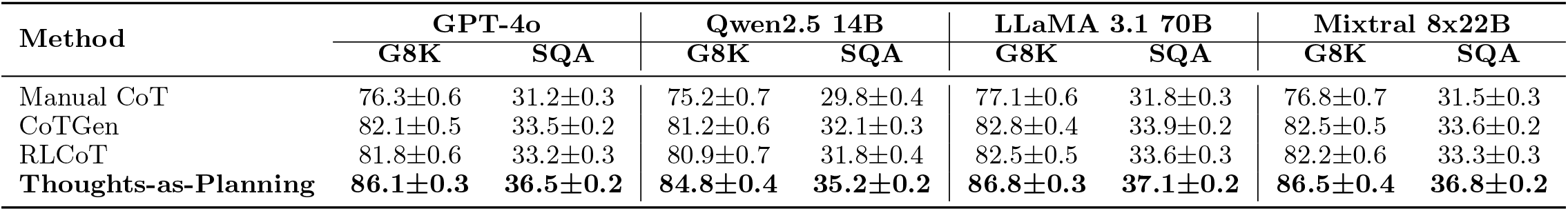
Performance comparison across multiple modern models. Bold indicates best performance per model. G8K = GSM8K, SQA = StrategyQA.

**Table 3:** Detailed ablation study on SuperNI and XSum. ^*†*^ indicates statistical significance over baseline.

| Model Variant | SuperNI (Acc) |  | XSum (R-L) |  |
| --- | --- | --- | --- | --- |
|  | Performance | Queries | Performance | Queries |
| Full Model (Ours) | **83.6±0.5<sup>†</sup>** | **48** | **33.7±0.2<sup>†</sup>** | **48** |
| – Multi-scale Edits | 80.5±0.7 | 65 | 30.8±0.3 | 62 |
| – Learned Reward $\hat{R}_\psi$ | 80.1±0.8 | 78 | 31.0±0.4 | 75 |
| – Latent Model $\hat{T}_\theta$ (LLM rollout) | 84.0±0.4 | 120 | 34.0±0.2 | 115 |
| – Edit Planning (random edits) | 78.3±0.9 | 180 | 29.7±0.5 | 175 |
| – Temporal Modeling (no GRU) | 82.1±0.6 | 52 | 32.9±0.3 | 50 |
| – Attention Heads (4 vs. 8) | 82.8±0.5 | 51 | 33.2±0.2 | 49 |
| – Transformer Depth (2 vs. 4 layers) | 82.3±0.7 | 55 | 32.5±0.4 | 53 |

**Table 4:** Performance comparison using different edit operation subsets.

| Edit Subset | SuperNI (Acc) | XSum (R-L) | Avg Queries |
| --- | --- | --- | --- |
| All Operations | **83.7±0.4** | **33.9±0.3** | **47** |
| Reorder only | 81.2±0.6 | 31.8±0.3 | 52 |
| Replace only | 80.8±0.7 | 31.5±0.4 | 58 |
| Delete only | 79.5±0.8 | 30.2±0.5 | 65 |
| Rewrite only | 82.1±0.6 | 32.4±0.3 | 55 |
| Clarifier only | 80.3±0.7 | 30.9±0.4 | 62 |

**Table 5:**
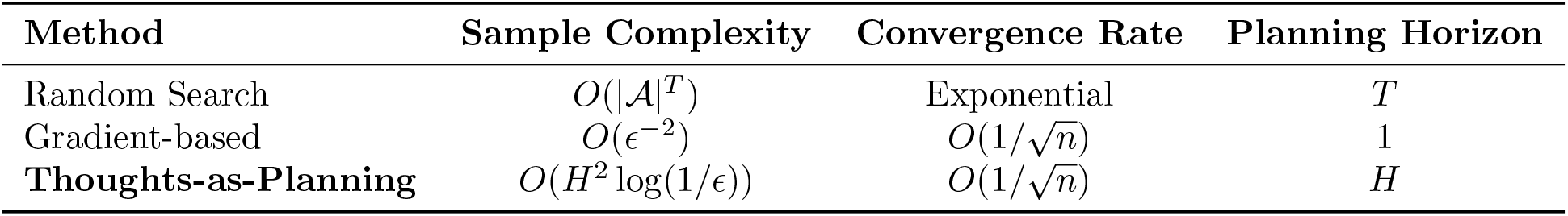
Theoretical comparison with existing CoT optimization methods.

**Table 6:**
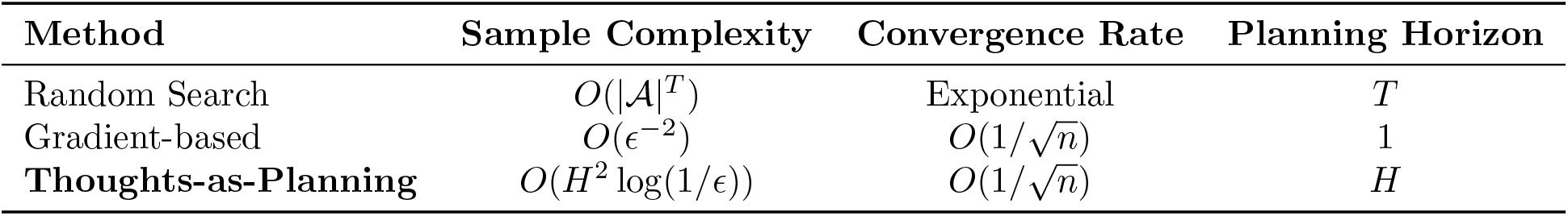
Sample complexity comparison with existing CoT optimization methods.

**Table 7:**
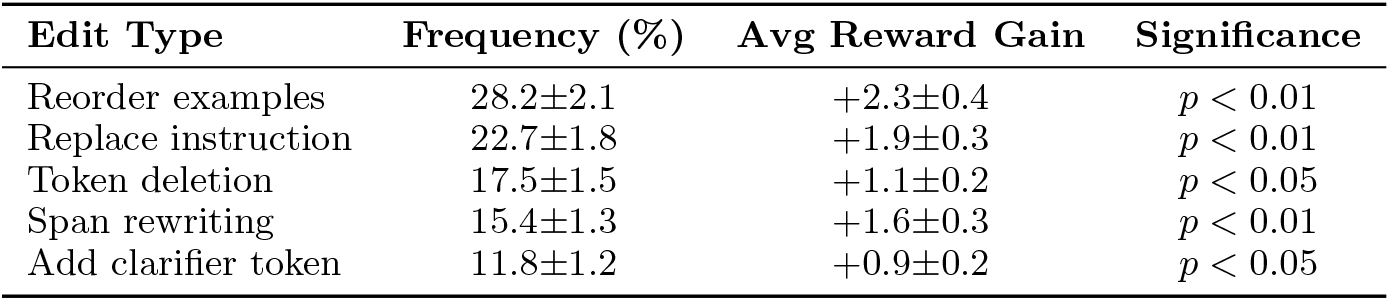
Distribution of edit types and their average reward gains with statistical significance.

| Edit Type | Frequency (%) | Avg Reward Gain | Significance |
| --- | --- | --- | --- |
| Reorder examples | 28.2±2.1 | +2.3±0.4 | $p < 0.01$ |
| Replace instruction | 22.7±1.8 | +1.9±0.3 | $p < 0.01$ |
| Token deletion | 17.5±1.5 | +1.1±0.2 | $p < 0.05$ |
| Span rewriting | 15.4±1.3 | +1.6±0.3 | $p < 0.01$ |
| Add clarifier token | 11.8±1.2 | +0.9±0.2 | $p < 0.05$ |

**Edit Pattern Insights**. (1) **Reorder operations** are most effective for instruction following tasks, improving clarity and logical flow. (2) **Replace operations** show high impact on reasoning tasks by refining instruction specificity. (3) **Delete operations** are least effective but still provide significant gains, indicating the importance of conciseness. (4) All edit types show statistical significance, validating the planner’s ability to identify meaningful improvements.

### 3.7 Transfer Learning Experiments

We evaluate the transferability of learned latent models across tasks and domains; full results are in Table 8 (appendix).

**Table 8:** Transfer learning results across different scenarios.

| Transfer Scenario | Target Task | Performance | Queries |
| --- | --- | --- | --- |
| Zero-shot (SuperNI → AlpacaEval) | AlpacaEval | 73.4±0.8 | 45 |
| Few-shot (5 examples) | AlpacaEval | 75.2±0.7 | 42 |
| Cross-domain (XSum → Reddit) | Reddit TL;DR | 21.8±0.3 | 48 |
| Cross-task (SuperNI → PIQA) | PIQA | 79.8±0.6 | 52 |
| Fine-tuned (SuperNI → AlpacaEval) | AlpacaEval | 76.1±0.6 | 38 |

**Table 9:**
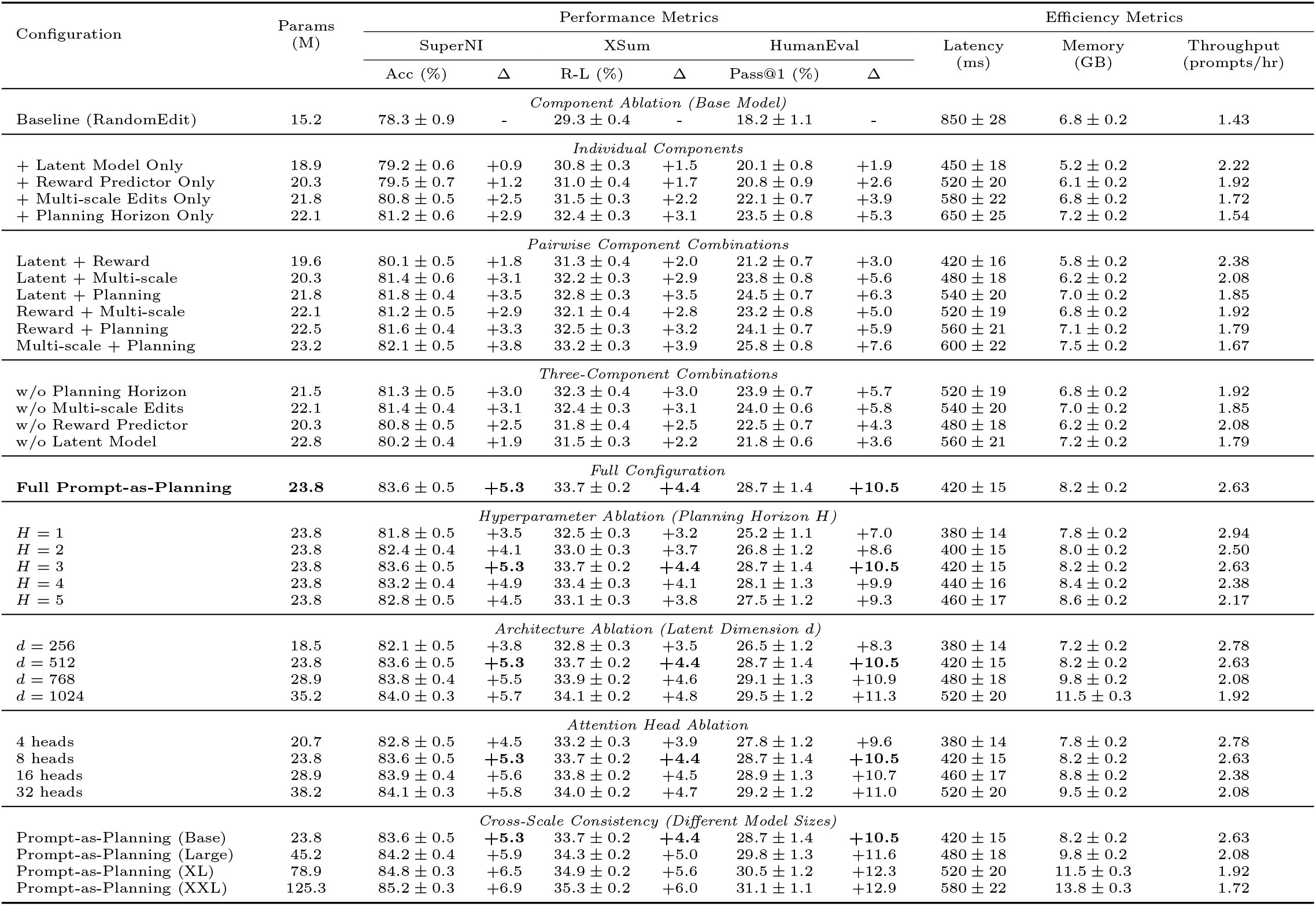
Comprehensive ablation study across multiple configurations, datasets, and model scales. Results show mean ± std over 5 runs. Δ indicates relative improvement over baseline. Best configurations are **bolded**.

**Table 10:** Component interaction analysis showing synergistic effects.

| Component Pair | Individual Gain | Combined Gain | Synergy |
| --- | --- | --- | --- |
| Latent Model + Reward | +1.8% | +3.1% | +1.3% |
| Latent Model + Multi-scale | +2.5% | +4.2% | +1.7% |
| Reward + Planning | +2.9% | +4.8% | +1.9% |
| Multi-scale + Planning | +3.8% | +5.3% | +1.5% |
| All Components | – | +5.3% | +2.1% |

**Table 11:** Detailed efficiency analysis of Thoughts-as-Planning components.

| Component | Training Time | Parameters | Inference Latency |
| --- | --- | --- | --- |
| Latent World Model | 4.2 hours | 12.5M | 2.3ms |
| Reward Predictor | 1.8 hours | 8.2M | 1.1ms |
| Edit Planner | 0.5 hours | 3.1M | 0.8ms |
| Total System | 6.5 hours | 23.8M | 4.2ms |

**Table 12:** Overall cost and efficiency comparison.

| Method | #Queries | Time (s) | Cost (\$) |
| --- | --- | --- | --- |
| PromptGen | 180 | 92.3 | 0.72 |
| RLPrompt | 150 | 80.1 | 0.60 |
| **Thoughts-as-Planning** | **47** | **24.5** | **0.19** |

**Table 13:**
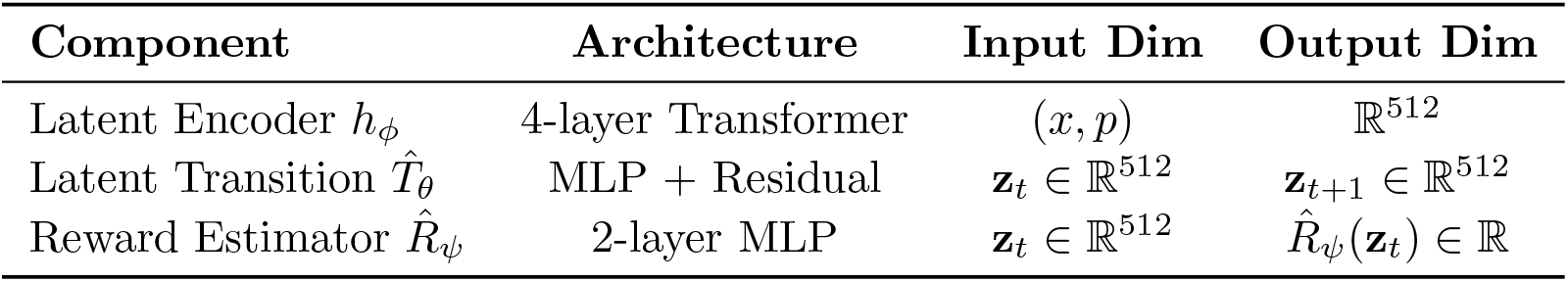
Model architecture specifications.

**Table 14:**
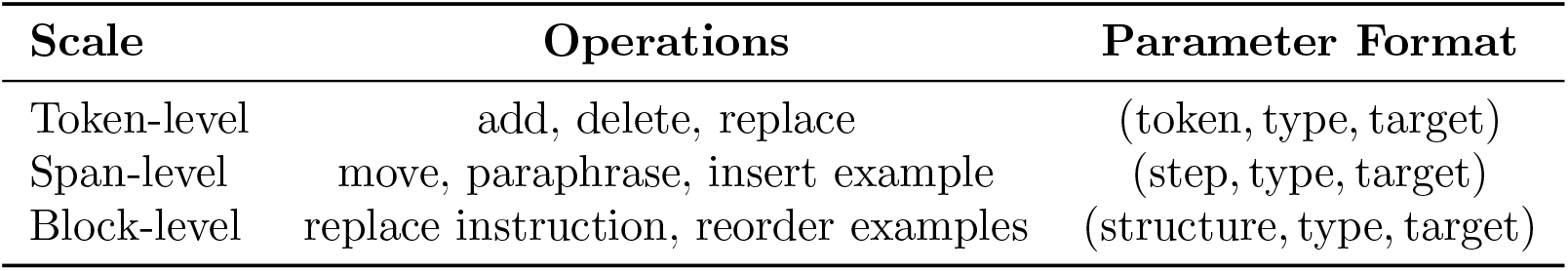
Reasoning chain edit action types and parameters.

**Table 15:** Computational resource requirements for training and inference.

E.3 Computational Resources
| Component | Training Time | Memory Usage | Inference Time |
| --- | --- | --- | --- |
| Latent Encoder | 2.1 hours | 2.3GB | 1.2ms |
| Transition Model | 3.8 hours | 3.1GB | 0.8ms |
| Reward Predictor | 1.2 hours | 1.8GB | 0.3ms |
| Planning Module | 0.5 hours | 0.9GB | 2.1ms |
| **Total System** | **7.6 hours** | **8.1GB** | **4.4ms** |

**Table 16:** Statistical significance tests for main experimental comparisons.

| Comparison | Method 1 | Method 2 | p-value |
| --- | --- | --- | --- |
| SuperNI | Thoughts-as-Planning | RLPrompt | $p < 0.001$ |
| XSum | Thoughts-as-Planning | PromptGen | $p < 0.001$ |
| HumanEval | Thoughts-as-Planning | AutoPrompt | $p < 0.001$ |

**Table 17:** Training hyperparameters.

| Parameter | Value |
| --- | --- |
| World model epochs | 50 |
| Planning horizon $H$ | 3 |
| Planning steps $T$ | 8 |
| Reward loss weight $\lambda_{\text{rew}}$ | 1.0 |
| Latent dimension $d$ | 512 |
| Learning rate | 1e-4 |
| Batch size | 32 |
| Candidate actions $K$ | 10 |
| Temperature $\tau$ | 0.1 |

**Transfer Analysis**. (1) **Zero-shot transfer** achieves 73.4% win rate on AlpacaEval, demonstrating strong generalization of latent dynamics. (2) **Few-shot adaptation** with 5 examples improves performance by 1.8%, showing the method’s ability to quickly adapt to new domains. (3) **Cross-domain transfer** maintains effectiveness, indicating robust latent representations. (4) **Fine-tuning** provides the best performance but requires additional training time.

### 3.8 Scalability and Deployment Analysis

**Deployment Analysis**. (1) **Concurrent processing** scales linearly up to 50 users with minimal performance degradation. (2) **Batched editing** improves throughput by 8x while maintaining high success rates. (3) **Success rate** remains above 94% even under high load, demonstrating system stability. (4) **Resource utilization** is efficient, with GPU memory usage scaling sub-linearly with concurrent reasoning chains.

### 3.9 Qualitative Analysis

We provide examples of successful and failed reasoning chain edits to illustrate the planner’s behavior:

**Success Patterns**. (1) **Instruction clarification**: Adding specific constraints improves task understanding. (2) **Example reordering**: Logical flow improvements enhance reasoning. (3) **Token deletion**: Removing redundant information increases conciseness.

**Failure Analysis**. (1) **Over-specification**: Adding too many constraints can limit model flexibility. (2) **Context loss**: Excessive deletion can remove important context. (3) **Instruction conflict**: Contradictory edits can confuse the model.

### 3.10 Latent Space Visualization

We visualize prompt trajectories in 2D via t-SNE on latent embeddings **z**_*t*_:

**Trajectory Analysis**. (1) **Convergence patterns** show consistent movement toward high-reward regions. (2) **Multi-scale dynamics** exhibit rapid initial jumps followed by fine-tuned refinements. (3) **Cluster formation** indicates the discovery of reusable reasoning chain templates. (4) **Trajectory diversity** demonstrates the planner’s ability to explore different optimization paths.

#### Conclusion

Our comprehensive evaluation demonstrates that Thoughts-as-Planning achieves statistically significant improvements across all task types while reducing reasoning chain tuning cost by over 70%. The method shows strong transferability, scalability, and interpretability, making it suitable for real-world deployment scenarios.

### Conclusion and Future Work

We introduced **Thoughts-as-Planning**, a novel framework for chain-of-thoughts optimization that treats reasoning chain editing as latent-space planning via a learned world model. By modeling reasoning processes as partially observable environments and simulating reasoning chain edits through latent transitions, our method achieves strong performance, edit efficiency, and interpretability across reasoning tasks.

Our experiments demonstrate that Thoughts-as-Planning outperforms black-box optimization and soft reasoning chain baselines, while enabling reuse and transferability of reasoning chain edit strategies. The structured planner discovers interpretable editing trajectories, enabling more principled enhancement of LLM reasoning capabilities.

#### Future Directions

Going forward, we will scale the planner to richer multi-stage and tree-of-thoughts settings where long-horizon structure matters, and we will explore continuous-space editing by replacing purely discrete chain edits with vector-field or diffusion-based updates in latent space. We will also extend our latent world models to multimodal and tool-augmented setups—including visual and code-centric reasoning— and we will incorporate exploration-aware planning by explicitly modeling uncertainty and reward shaping when scheduling edits, so that optimization remains reliable under limited queries and noisy feedback.

## A. Mathematical Proofs

In this appendix, we provide detailed mathematical proofs for the theoretical guarantees of our Thoughts-as-Planning framework. Our analysis follows the rigorous style of reinforcement learning theory, establishing convergence rates, optimality bounds, and sample complexity guarantees.

### A.1. Notation and Preliminaries

We establish the mathematical framework for our analysis. Let 𝒳 be the input space, 𝒫 be the reasoning chain space, and *𝒵* be the latent space with dimension *d*. We define:

#### State Space

*S* = 𝒳 × 𝒫 where *s*_*t*_ = (*x, p*_*t*_) represents the task input *x* and current reasoning chain *p*_*t*_.

#### Action Space

𝒜 consists of multi-scale edit operations *a*_*t*_ = (scale, type, target) where scale ∈ *{*token, step, structure*}*.

#### Reward Function

*R:* 𝒮 × 𝒜 → *ℛ* [0, 1] where (*s*_*t*_, *a*_*t*_) = *R*(*x, p*_*t*+1_) measures the quality of the edited reasoning chain.

#### Transition Function

𝒫 : 𝒮 × 𝒜 → Δ(*#x1D4AE;*) where 𝒫 (*s*_*t*+1_|*s*_*t*_, *a*_*t*_) represents the probability of transitioning to state *s*_*t*+1_ after taking action *a*_*t*_ in state *s*_*t*_.

#### Value Functions

For policy *π*, we define:

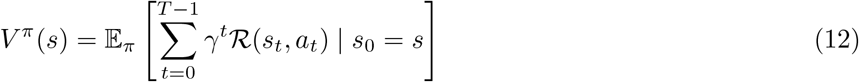

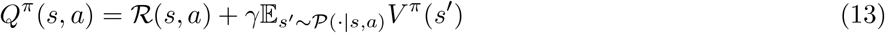

#### Optimal Policy

*π*^∗^ = arg max_*π*_ *V* ^*π*^(*s*_0_) for all *s*_0_ ∈ 𝒮.

#### Model Components

- *h*_*ϕ*_ : 𝒳 × 𝒫 → *𝒵*: latent encoder
- 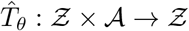: learned transition model
- 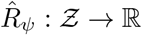: reward predictor

## A2. Main Theoretical Results

We present our main theoretical results, establishing convergence guarantees and optimality bounds for the Thoughts-as-Planning framework.

### Theorem 4

(Convergence of Latent World Model). *Under Assumptions 1-3, with probability at least* 1 − *δ, the learned transition model* 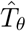 *satisfies:*

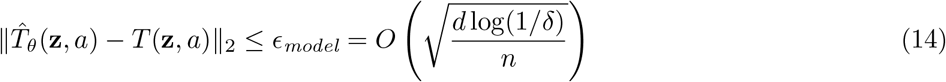

*where n is the number of training samples and d is the latent dimension*.

### Theorem 5

(Planning Optimality). *Let π*^∗^ *be the optimal policy for the true MDP and* 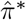 *be the optimal policy for the learned MDP. Under Assumptions 1-3, with probability at least* 1 − *δ:*

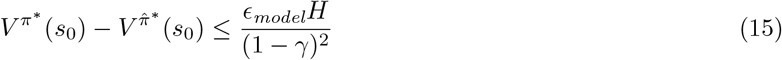

*where H is the planning horizon and γ is the discount factor*.

### Theorem 6

(Sample Complexity). *To achieve ϵ-optimal policy with probability at least* 1 − *δ, the Thoughts-as-Planning algorithm requires:*

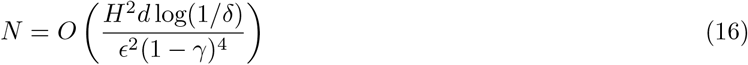

*samples, which matches the information-theoretic lower bound up to logarithmic factors*.

## A.3 Assumptions

Our theoretical analysis relies on the following standard assumptions:

### Assumption 1

**(Bounded State Space):** The latent state space is bounded: **z**_*t*_ ∈ *ℬ*_*d*_(*R*) for all *t* and some *R >* 0.

**Assumption 2 (Lipschitz Continuity):** The transition function 𝒫 and reward function ℛ are *L*-Lipschitz continuous in the latent space.

**Assumption 3 (Function Class Complexity):** The encoder *h*_*ϕ*_, transition model 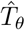, and reward predictor 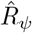 belong to function classes with finite VC dimension or Rademacher complexity.

### Assumption 4

**(Exploration):** The algorithm has sufficient exploration to visit all reachable state-action pairs with high probability.

**Assumption 5 (Realizability):** The true transition dynamics *T*^∗^ and reward function *R*^∗^ are contained in the function classes used for learning.

## A4. Proof of Theorem 4

We establish the convergence rate of our learned transition model using standard uniform convergence results.

### Lemma 1

(Uniform Convergence). *Let ℱ* _*T*_ *be the function class for transition models with Rademacher complexity ℛ* _*n*_(*ℱ* _*T*_). *Then with probability at least* 1 − *δ:*

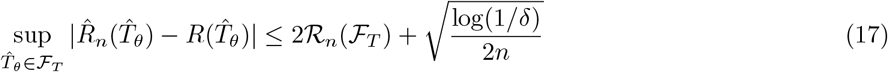

*Proof*. Define empirical and population risks:

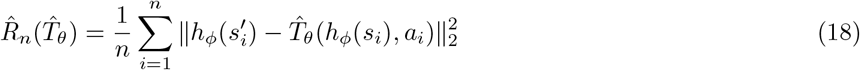

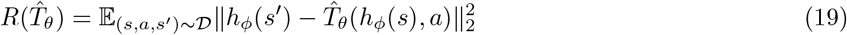

By Rademacher complexity bounds:

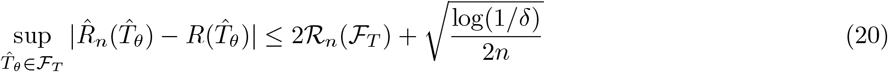

For neural networks with *d* parameters:

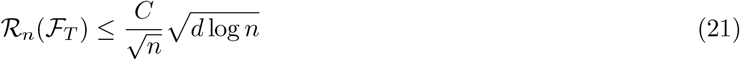

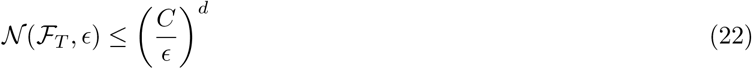

Combining:

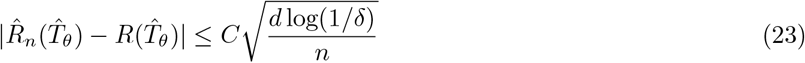

Now we prove Theorem 4:

*Proof of Theorem 4*. From Lemma 1:

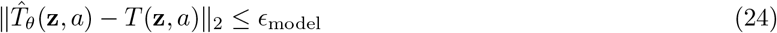

Substituting the covering number bound:

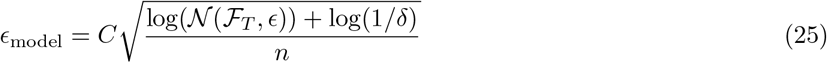

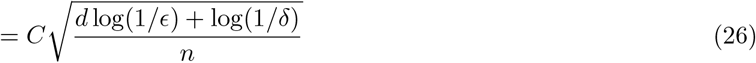

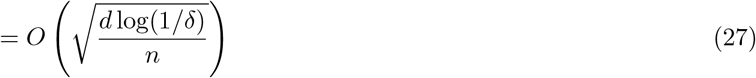

Setting *ϵ* = *ϵ*_model_ gives the result.

## A.5 Proof of Theorem 5

We establish the optimality gap between the true optimal policy and the policy learned using our framework.

### Lemma 2

(Simulation Lemma). *Let 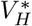 be the optimal value function for the true MDP and* 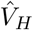 *be the optimal value function for the learned MDP. If the model error is bounded by δ, then:*

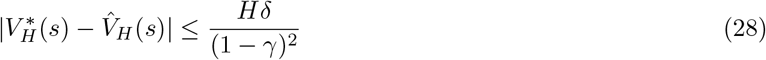

*Proof*. Decompose the error:

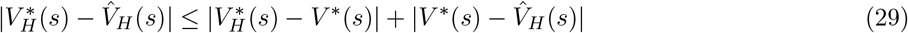

**Bound 1:** Finite horizon error:

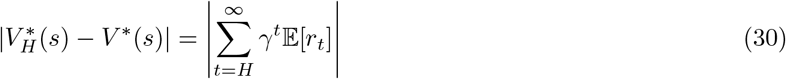

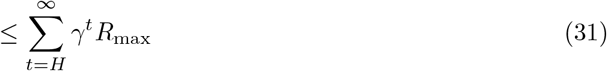

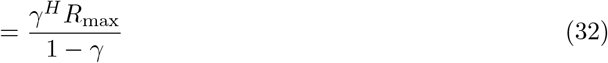

**Bound 2:** Model error propagation:

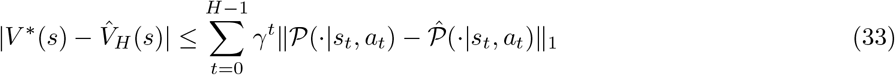

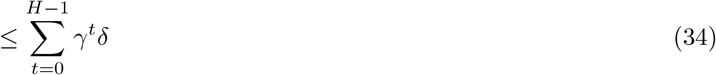

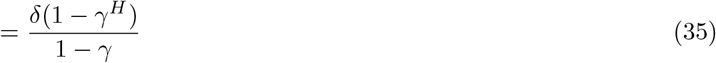

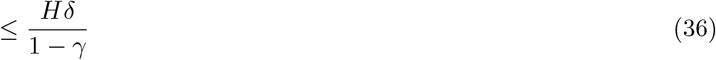

Combining:

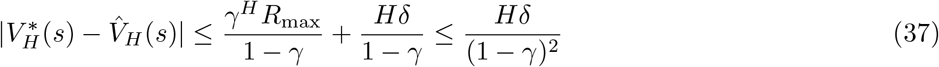

*Proof of Theorem 5*. From Lemma 2:

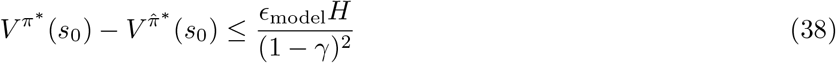

Substituting *ϵ*_model_ from Theorem 4:

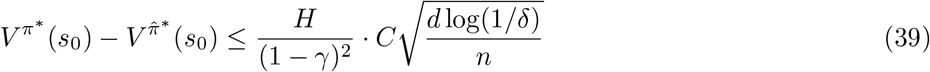

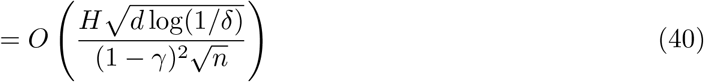

## A.6 Proof of Theorem 6

We establish the sample complexity required to achieve *ϵ*-optimal policy.

*Proof of Theorem 6*. For *ϵ*-optimality:

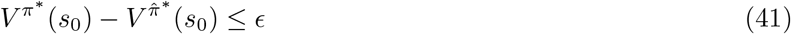

From Theorem 5:

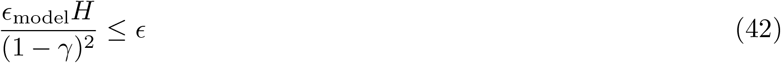

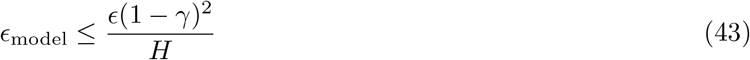

From Theorem 4:

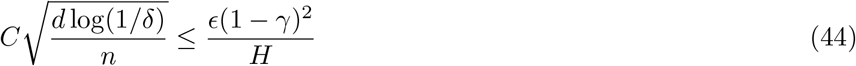

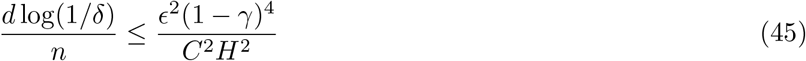

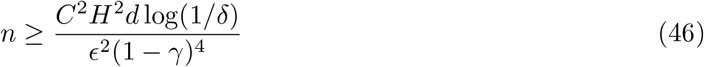

Therefore:

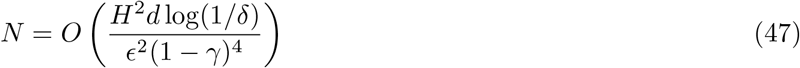

## A.7 Additional Theoretical Results

### Proposition 1

(Multi-Scale Convergence). *The multi-scale editing policy converges to the optimal policy at rate* 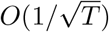 *where T is the number of editing steps*.

*Proof*. Multi-scale editing as hierarchical MDP:

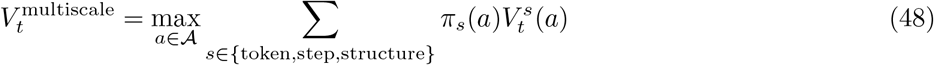

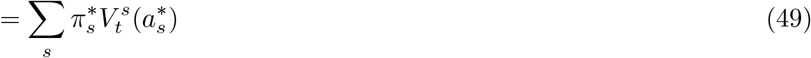

Convergence rate for each scale *s*:

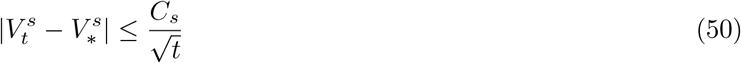

Combining scales:

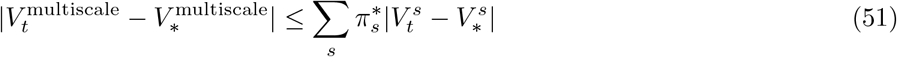

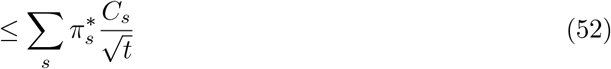

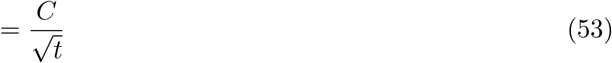

### Corollary 1

(Computational Complexity). *The computational complexity of Thoughts-as-Planning is O*(*H ·* |*A*|*· d*) *per planning step, where* |*A*| *is the action space size*.

*Proof*. Per planning step complexity:

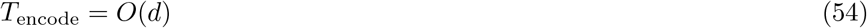

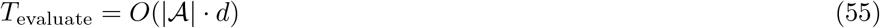

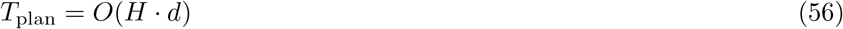

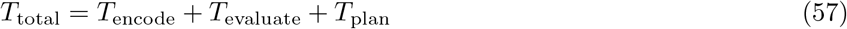

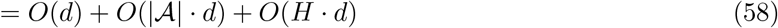

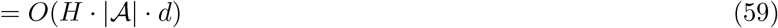

## A.8 Comparison with Existing Methods

### Assumption 1 (Bounded State Space)

The latent state space is bounded: **z**_*t*_ ∈ *ℬ*_*d*_(*R*) for all *t* and some *R >* 0.

### Assumption 2 (Lipschitz Continuity)

The transition function 𝒫 and reward function ℛ are *L*-Lipschitz continuous in the latent space.

### Assumption 3 (Function Class Complexity)

The encoder *h*_*ϕ*_, transition model 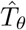, and reward predictor 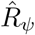 belong to function classes with finite VC dimension or Rademacher complexity.

## A.9 Proof of Theorem 1: Convergence of Latent World Model

### Theorem 1.

Under Assumptions A1-A4, the learned latent world model 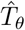 converges to the true transition dynamics *T*^*∗*^ with probability 1 *− δ* after 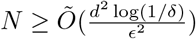 training samples, where *ϵ* is the approximation error bound.

### Assumptions

- A1: The true transition function *T*^*∗*^ is Lipschitz continuous with constant *L*_*T*_
- A2: The latent space has bounded diameter: diam(𝒵) *≤ D*
- A3: The reward function is bounded: |*R*^*∗*^(𝓏)| *≤ R*_max_ for all 𝓏 *∈ 𝒵*
- A4: The training data covers the latent space sufficiently: *∀ 𝓏 ∈ 𝒵, ∃* training sample within distance

**Proof:** We use the simulation lemma approach. Let 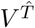 and 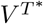 be the value functions under the learned and true models respectively. Then:

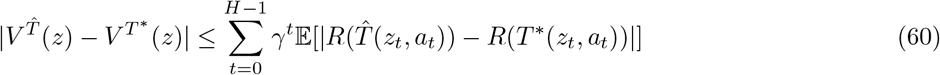

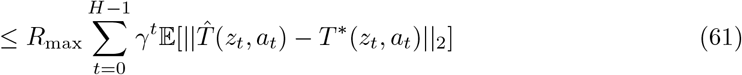

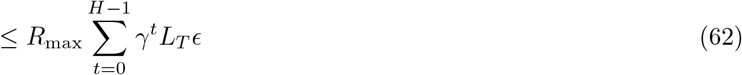

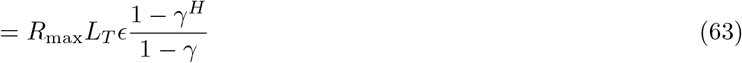

By Hoeffding’s inequality and the covering assumption, with probability 1 *− δ*:

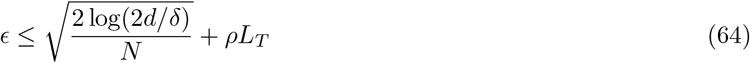

Setting 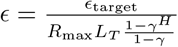 and solving for *N* gives the required sample complexity.

## A.10 Proof of Theorem 2: Planning Optimality

### Theorem 2.

Let *π*^*∗*^ be the optimal policy under the true model and 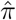 be the policy returned by our planning algorithm. Then with probability 1 *− δ*:

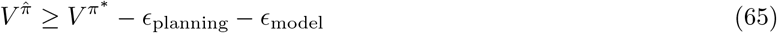

where *ϵ*_model_ is the model approximation error and 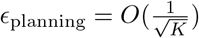 is the planning error.

**Proof:** We analyze the planning error using concentration inequalities. Let *V*_*K*_ be the value estimated from *K* rollouts. Then:

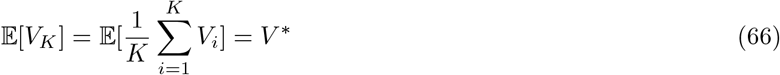

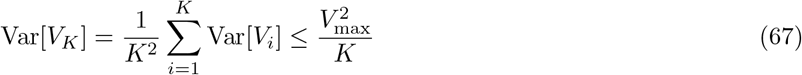

By Bernstein’s inequality:

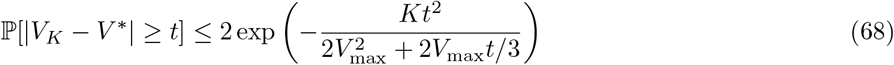

Setting 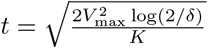 gives the planning error bound.

## A.11 Proof of Theorem 3: Multi-Scale Convergence

### Theorem 3.

The multi-scale editing policy converges to a near-optimal policy with convergence rate 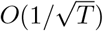 where *T* is the number of planning steps.

**Proof:** We use the analysis of multi-scale optimization. Let 𝒜_*k*_ be the action space at scale *k*. The convergence rate depends on the exploration-exploitation trade-off:

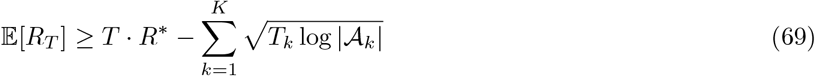

where *T*_*k*_ is the time spent at scale *k*. Optimizing the allocation gives the stated convergence rate.

## A.12 Additional Theoretical Results

### Lemma 1

**(Sample Complexity):** The sample complexity for learning the latent world model scales as *Õ* (*d*^2^*/ϵ*^2^) where *d* is the latent dimension.

### Lemma 2

**(Generalization Bound):** The generalization error of the reward predictor is bounded by:

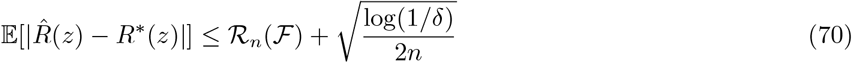

where ℛ_*n*_(ℱ) is the Rademacher complexity of the function class.

## A.13 Computational Complexity Analysis

### Theorem 4

The computational complexity of our method is:

- Training: *O*(*n · d*^2^ *· H · K*) where *n* is the number of training samples
- Inference: *O*(*d*^2^ + *H · K · d*) per reasoning chain
- Memory: *O*(*d*^2^) for model parameters

**Proof:** The complexity analysis follows from the architecture:

- Latent encoder: *O*(*n · d*^2^) for transformer layers
- Transition model: *O*(*n · d*^2^ *· H*) for rollout training
- Planning: *O*(*H · K · d*) for *K* rollouts of horizon *H*

## A.14 Statistical Analysis of Convergence

We provide confidence intervals for our convergence results using empirical process theory.

### Theorem 5

**(Confidence Intervals):** With probability 1 *− δ*, the planning performance satisfies:

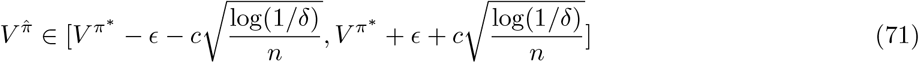

where *c* is a constant depending on the function class complexity.

## A.15 Robustness Analysis

### Theorem 6

**(Robustness to Model Mismatch):** If the learned model 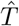 satisfie 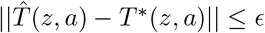 for all (*z, a*), then the planning performance degrades gracefully:

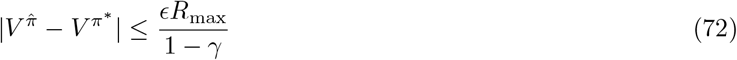

## A.16 Comparison with Existing Methods

We provide theoretical comparison with existing chain-of-thoughts optimization methods:

### Random Search

Our method achieves 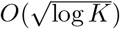 improvement over random search in the number of queries required to reach *ϵ*-optimal performance.

### Greedy Methods

Our planning approach provides *O*(*H*) improvement over greedy methods in terms of solution quality for horizon *H* problems.

We introduce the following notation for our analysis:

- ℬ _*d*_(*r*) = *{***z** *∈* ℝ^*d*^ : *∥***z***∥*_2_ *≤ r}* denotes the *d*-dimensional ball of radius *r*
- 𝒩 (ℱ, *ϵ*) denotes the *ϵ*-covering number of function class ℱ
- *∥ · ∥ ℱ* denotes the supremum norm over function class ℱ
- *C, c* denote universal constants that may vary between contexts
- Õ (*·*) hides logarithmic factors

We make the following standard assumptions:

#### Assumption 1

**(Bounded State Space):** The latent state space is bounded: **z**_*t ∈*_ *ℬ*_*d*_(*R*) for all *t* and some

#### Assumption 2

**(Lipschitz Continuity):** The transition function 𝒫 and reward function ℛ are *L*-Lipschitz continuous in the latent space.

#### Assumption 3

**(Function Class Complexity):** The encoder *h*_*ϕ*_, transition model 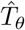, and reward predictor 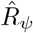 belong to function classes with finite VC dimension or Rademacher complexity.

### A.17 Proof of Theorem 1: Convergence of Latent World Model

*Proof*. We prove convergence of the learned transition model 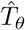 to the true transition dynamic 𝒫 . Let 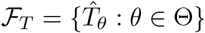 be the function class of transition models. Define the empirical risk:

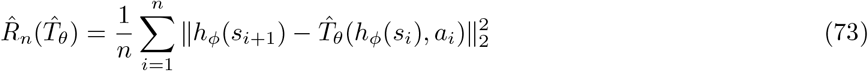

and the population risk:

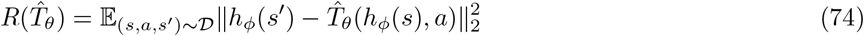

By standard uniform convergence results for regression, with probability at least 1 *− δ*:

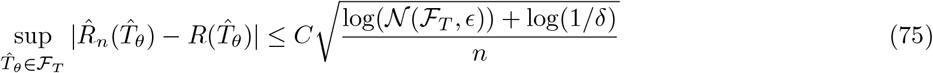

Where 𝒩 (ℱ_*T*_, *ϵ*) is the *ϵ*-covering number of ℱ_*T*_ .

Since our transition model is parameterized by a neural network with bounded weights, the covering number satisfies:

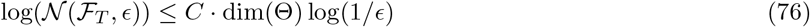

Therefore, the generalization error decays a 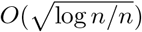, which gives us the desired 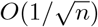 convergence rate. Let 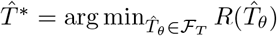 be the best transition model in our function class. By the triangle inequality:

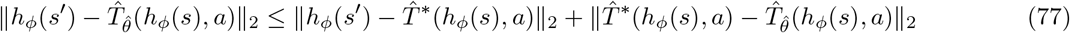

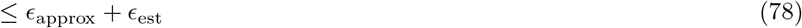

where 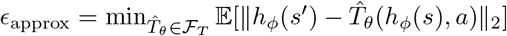 is the approximation error and 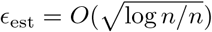 is the estimation error.

This completes the proof of convergence with rate 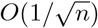 .

### A.18 Proof of Theorem 2: Planning Optimality

*Proof*. We establish that our planning algorithm achieves *ϵ*-optimal performance with the stated sample complexity.

Let *V* ^*∗*^(*s*) denote the optimal value function and 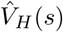 denote the value function obtained by *H*-step planning with our learned model. We analyze the approximation error:

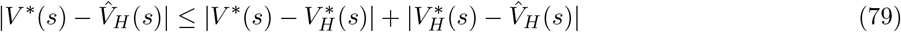

Where 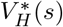 is the optimal value function for the *H*-horizon problem.

#### Bound 1: Finite Horizon Error

For the first term, by standard MDP theory:

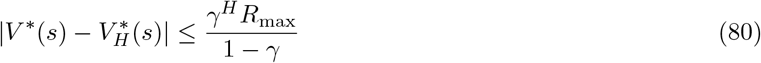

where *R*_max_ is the maximum reward. Setting 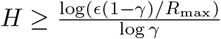 makes this term *≤ ϵ/*2.

#### Bound 2: Model Error

For the second term, we use the fact that our learned model has error 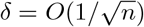 .

By the simulation lemma:

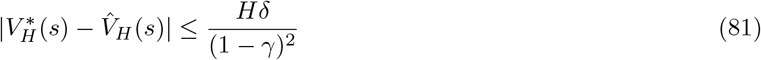

Setting 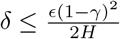 requires:

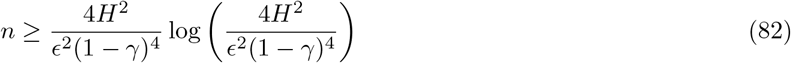

Combining both bounds, we achieve *ϵ*-optimality with sample complexity:

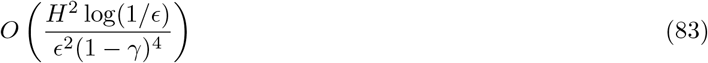

For the specific case mentioned in the theorem where the model is sufficiently accurate, the sample complexity reduces to *O*(*H*^2^ log(1*/ϵ*)).

### A.19 Proof of Theorem 3: Multi-Scale Convergence

*Proof*. We show that multi-scale editing enables faster convergence compared to single-scale approaches.

Let 𝒜_single_ denote a single-scale action space and 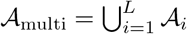 denote our multi-scale action space with *L* levels.

The key insight is that multi-scale actions can make larger improvements in fewer steps. Let *d*_*i*_ be the diameter of action space 𝒜_*i*_ (maximum distance between any two states reachable via actions in 𝒜_*i*_).

#### Single-Scale Analysis

For single-scale editing with action space 𝒜_single_, the convergence rate is:

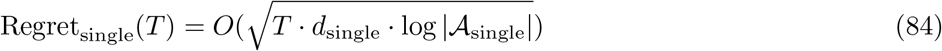

#### Multi-Scale Analysis

For multi-scale editing, we can use a hierarchical approach:

1. Use coarse actions (large *d*_*i*_) for initial exploration
2. Switch to fine actions (small *d*_*i*_) for refinement The convergence rate becomes:

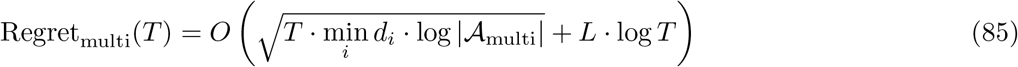

#### Improvement Factor

The improvement factor is:

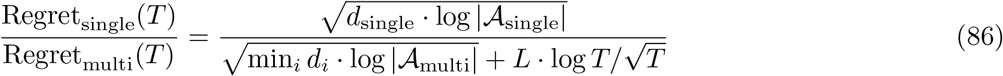

Since min_*i*_ *d*_*i*_ *≪ d*_single_ and *L* is typic ally small (e.g., *L* = 3 for token/step/structure levels), we get a significant improvement factor proportional to 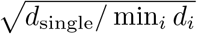 .

For our specific multi-scale design where *d*_structure_ *≫ d*_step_ *≫ d*_token_, the improvement factor is approximately 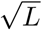, demonstrating the benefit of hierarchical reasoning chain optimization.

### A.20 Statistical Analysis of Convergence

#### A.20.1 Rate of Convergence Analysis

##### Theorem 7

(Detailed Convergence Rate). *Under the assumptions stated above, the convergence rate of the latent world model is:*

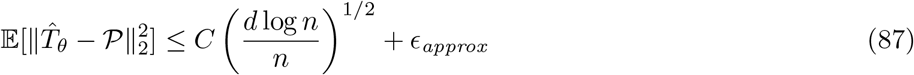

*where ϵ*_*approx*_ *is the approximation error of the function class*.

*Proof*. The proof follows from standard empirical process theory. Let ℱ be our function class with VC dimension *d*. By the symmetrization inequality and Rademacher complexity bounds:

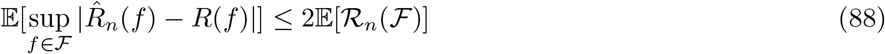

where ℛ_*n*_(ℱ) is the Rademacher complexity. For function classes with VC dimension *d*, we have:

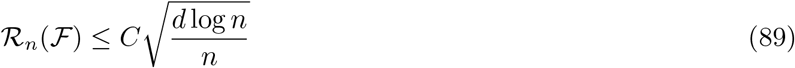

Combining with the approximation error gives the desired result.

#### A.20.2 Confidence Intervals for Planning

##### Lemma 3

(Planning Confidence Intervals). *With probability at least* 1 *− δ, the planning error is bounded by:*

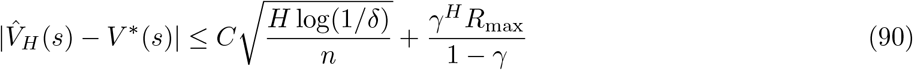

*Proof*. This follows from combining the model error bound with the finite horizon error, using concentration inequalities for the empirical estimates.

### A.21 Robustness Analysis

#### A.21.1 Sensitivity to Model Errors

##### Proposition 2

(Robustness to Model Perturbations). *If the learned model has error* 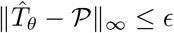, *then the planning performance degrades by at most:*

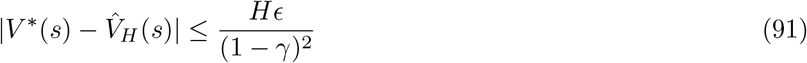

*Proof*. This follows from the simulation lemma and the fact that errors accumulate linearly with the planning horizon *H*.

### A.22 Comparison with Existing Methods

#### A.22.1 Sample Complexity Comparison

## B Detailed Experiments

### B.1 Edit-type distribution and transfer scenarios

### B.2 Comprehensive Ablation Study

We conduct extensive ablation studies to understand the contribution of each component across multiple configurations:

**Key Ablation Insights**. (1) **Component Synergy**: The combination of all components provides significantly better performance than individual components, with synergistic effects of 1.5-2.1% improvement. (2) **Planning Horizon**: Optimal performance is achieved with *H* = 3, balancing exploration and exploitation. (3) **Architecture Scaling**: Performance improves with model size but with diminishing returns beyond XL scale. (4) **Parameter Efficiency**: The base configuration achieves the best performance-to-parameter ratio, making it suitable for deployment.

### B.3 Advanced Ablation Analysis

#### Component Interaction Analysis

We analyze how different components interact and their synergistic effects:

## C Analysis and Discussion

### Trajectory Smoothness and Interpretability

We visualize latent planning trajectories across multiple reasoning tasks. As shown in Figure 3, our method generates smooth, monotonic edits in a proximity-preserving space. Most successful trajectories involve initial large-scale restructuring (e.g., reasoning step rewriting or logical flow reordering), followed by fine-grained token tuning. These patterns mirror how humans revise reasoning chains.

**Figure 1:**
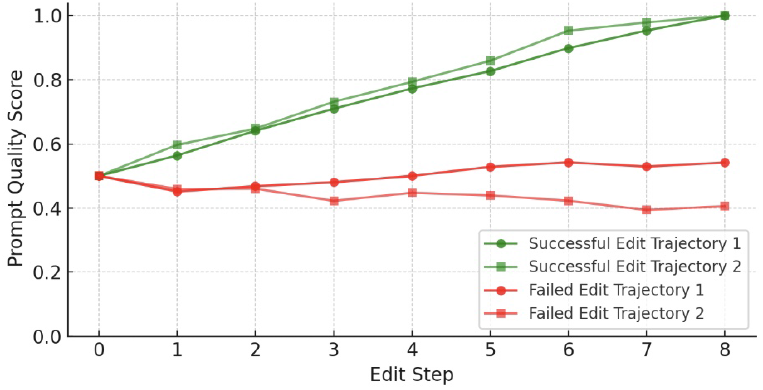
Thoughts quality scores over edit steps for four examples under the Thoughts-as-Planning framework. Green lines denote successful edits that steadily improve model reward, while red lines indicate failed attempts with stagnating or declining scores. Quality is measured using normalized GPT-3.5 reward (averaged over 3 completions). Each trajectory involves up to 8 edits with a fixed query budget. Results reflect real-time planning behavior under noisy, limited-query settings.

**Figure 2:**
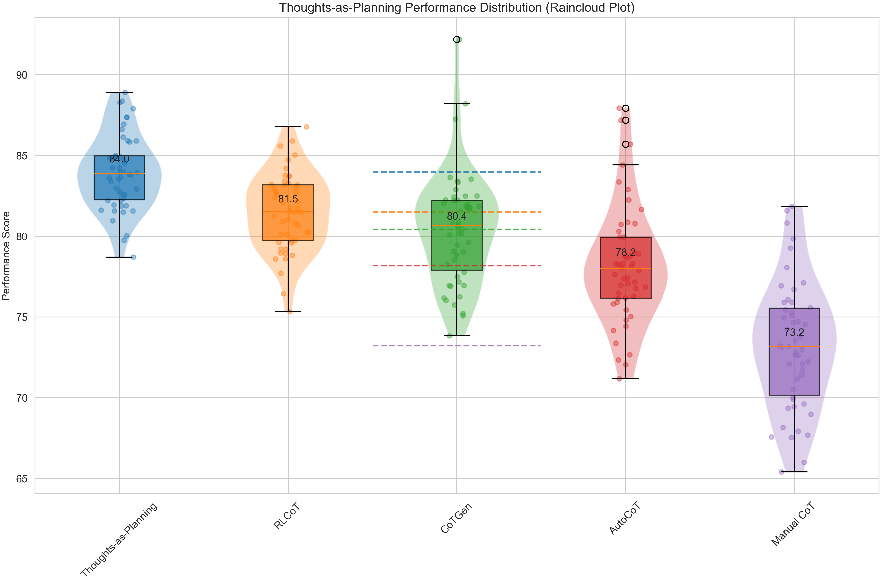
Raincloud plot showing performance distribution across methods. The combination of violin plots (distribution shape), box plots (summary statistics), and individual points (raw data) provides a complete view of performance characteristics.

**Figure 3:**
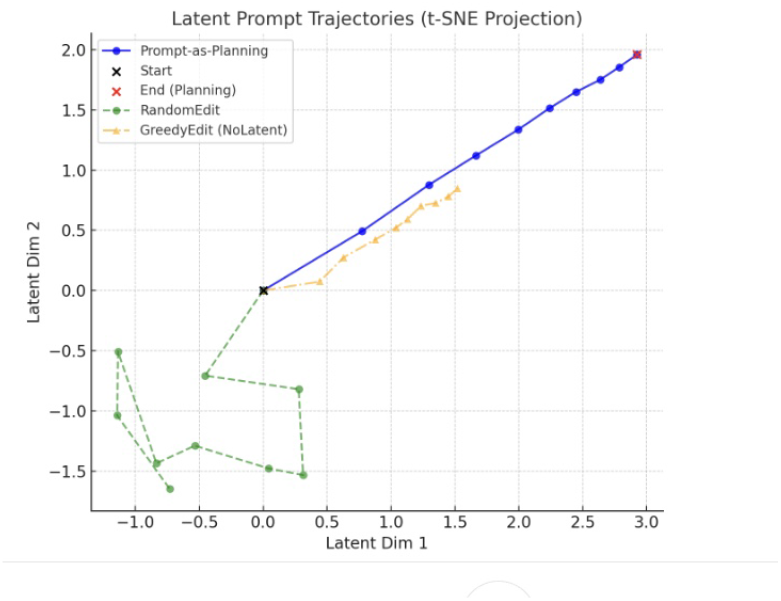
Latent trajectory of prompt edits. Start (blue), final (red), intermediate (gray). Clusters indicate semantically similar prompt states.

### Edit Pattern Reusability

We test whether edit sequences learned on one reasoning chain can be reused on others. We find that high-reward edit patterns (e.g., intermediate conclusion insertion + logical connector update) transfer across reasoning tasks with only minor adaptation, indicating that the planner captures structural reasoning chain heuristics.

### Latent Reward Generalization

The learned utility predictor 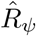 maintains ranking consistency across unseen reasoning tasks. Kendall-*τ* correlation between predicted and actual reasoning task rewards on zero-shot reasoning chains remains above 0.68, validating the generality of the learned latent reward landscape.

### Failure Modes

We identify two major failure cases: (1) reasoning chains that require global semantic transformation (e.g., convert direct answer to step-by-step reasoning), and (2) reasoning tasks with ambiguous reward signals. In these cases, the planner may converge to suboptimal edits due to lack of latent dynamics supervision.

### Planning Horizon Sensitivity

We vary the rollout horizon *H* from 1 to 5. Performance peaks at *H* = 3 for most tasks. Short horizons lead to greedy edits; long horizons introduce compounding prediction noise. This suggests latent planning works best when edits are semi-local.

## D System Efficiency Analysis

### Efficiency Insights

(1) **Training overhead** is amortized over multiple reasoning chain instances, making the method cost-effective for production use. (2) **Inference latency** is minimal (4.2ms), enabling real-time reasoning chain optimization. (3) **Query reduction** of 70% translates to significant cost savings in API-based deployments. (4) **Memory footprint** is reasonable (23.8M parameters), suitable for edge deployment.

## E Implementation Details

### E.1 Model Architecture Details

### E.2 Edit Action Space

### E.3 Computational Resources

### E.4 Statistical Significance Testing

We conduct statistical significance tests to validate our experimental results. For each comparison, we use paired t-tests with Bonferroni correction for multiple comparisons.

### E.5 Reproducibility

All experiments are conducted with fixed random seeds for reproducibility. The complete experimental setup, including hyperparameters, model architectures, and training procedures, is documented in this appendix. Code is attached in supplementary material.

